# Morphometric characters in the taxonomic recognition of the species of *Dolichopus pennatus* group (Diptera: Dolichopodidae)

**DOI:** 10.1101/2022.02.25.481977

**Authors:** M.A. Chursina, O.V. Selivanova

## Abstract

Sexually dimorphic structures (ornaments) on the legs are important diagnostic features for *Dolichopus* males, whereas females of those males are often morphologically similar and their species diagnostics is difficult. To assessing the relevance and feasibility of setting diagnostic characters for five sister species of the genus *Dolichopus* Latreille, 1796 (*D. argyrotarsis, D. lineaticornis, D. pennatus, D. popularis, D. subpennatus*) analysis of morphometric trait was used: comparison of wing shape by methods of geometric morphometry, as well as comparison of the ratios of the leg segments lengths. Comparative analysis of morphometric characters with molecular data made it possible to study the phylogenetic signal of male leg ornaments. It was shown for *Dolichopus* species, that modifications of the males’ middle legs occurred independently several times in the genus. In addition, the evolutionary pattern in the formation of similar ornaments was also associated with changes in the morphometric features of the legs and wings. Based on wing and legs morphometry, new diagnostic characters were proposed for females of the studied species group.

## Introduction

The genus *Dolichopus* Latreille, 1796 has an extremely high diversity with about 650 species [Grichanov, 2021] and is the largest genus of the Dolichopodidae family. Species of the genus are wide distributed and exhibit the greatest diversity in the Palaearctic region. Males often have characteristic ornaments such us flattened dorsoventrally or plumose segments of fore (*D. plumitarsis* Fallen, 1823), middle (*D. pennatus* Meigen, 1824) (Fig. 1) or hind tarsi (*D. remipes* Wahlberg, 1839), thickened hind tibia (*D. lepidus* Staeger, 1842), less frequently, enlarged arista (*D. jacutensis* Stackelberg, 192) or modification in wing coloration: for instance, males of *D. remipes* Wahlberg, 1839 have infuscated wings, and white spot at the apical part of wing is a diagnostic feature of males of *D. apicalis* Zetterstedt, 1849. Such interspecific variation suggests there has been significant evolutionary change in this trait, driven by sexual selection, but a robust phylogenetic hypothesis is required to study these changes. However, at the same time, females of the genus *Dolichopus* are difficult to diagnose using traditional taxonomic techniques and can often be discriminated only by legs color. However, as frequently noted, such characters are comparative, have low phylogenetic significance [Bernasconi et al., 2007b; Chursina, Grichanov, 2019], and their variation has never been examination in detail as a separate subject-matter in *Dolichopus* species.

**Fig. 1.**
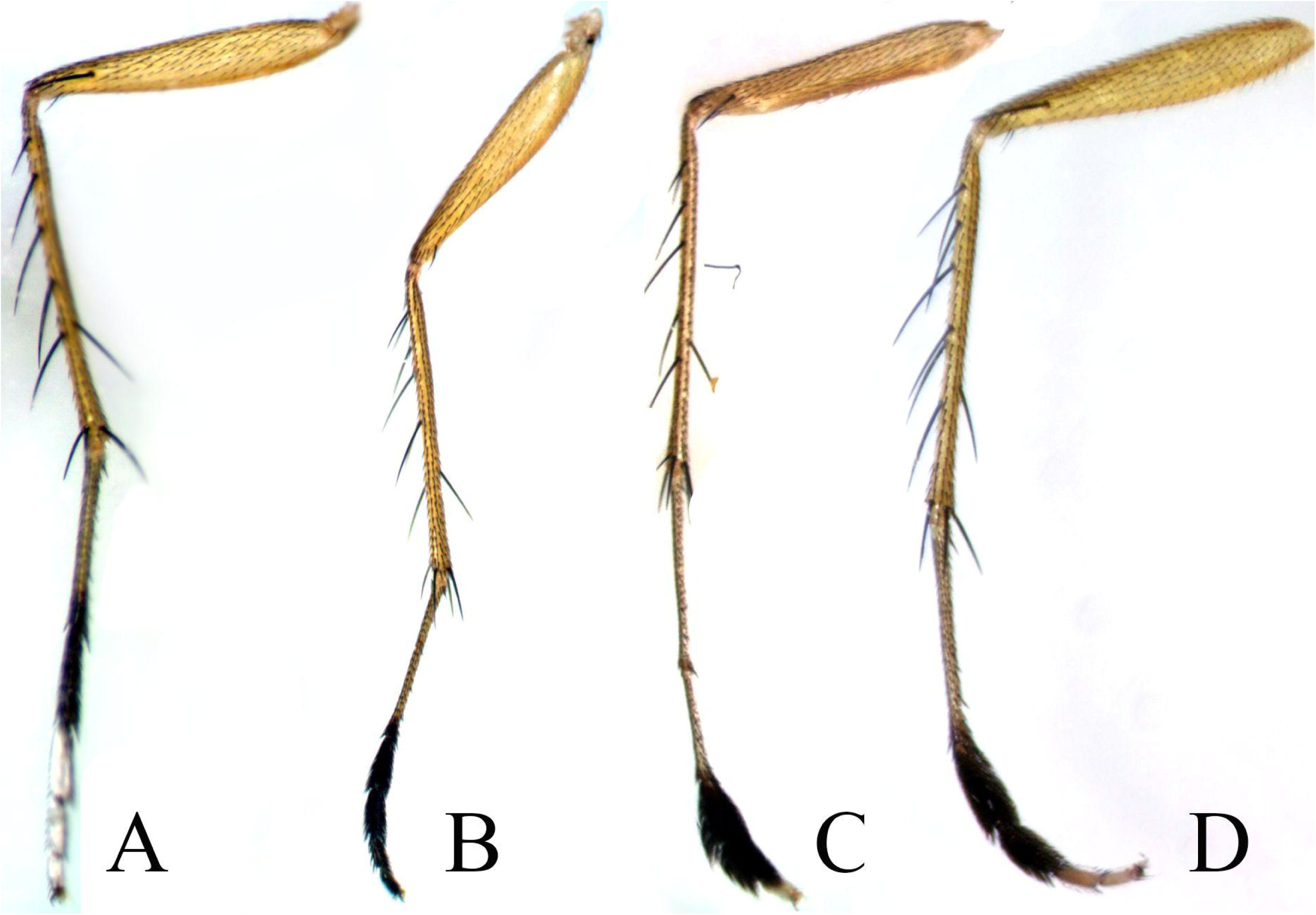
Ornaments on the middle legs of *Dolichopus* male: a – *D. argyrotarsis*; b – *D. pennatus*; c – *D. popularis*; d – *D. subpennatus*.

The geometric morphometric techniques analyzing represent a promising approach for discriminant between morphologically similar taxa [Pepinelli et al. 2013]. In present study, we examined differences in the morphometric characters between five *Dolichopus* species from sister group: *D. argyrotarsis* Wahlberg, 1850, *D. lineatocornis* Zetterstedt, 1843, *D. pennatus* Meigen, 1824, *D. popularis* Wiedemann, 1817, *D. subpennatus* d’Assis Fonseca, 1976.

Despite the fact, that Dolichopus species are widespread, the studied species are rare. They prefer moist biotopes: floodplain meadows, river and stream banks, swamps, deciduous forests. *D. argyrotarsis* distributed in Europe, including the European part of Russia [Negrobov et al., 2013]. The main distinguishing feature of the males of this species is enlarged and silver third, fourth and fifth segments of middle tarsi (fig. 1). The species have a Euro-Caucasian or Euro-Siberian range. The eastern boundaries of *D. pennatus* and *D. popularis* ranges include the Krasnoyarsk Territory (*D. popularis*) and Yakutia (*D. pennatus*). These two species is easily distinguished from other *Dolichopus* males by an enlarged segments of middle tarsi, but males of *D. pennatus* has enlarged and plumose second and third tarsi segments, whereas *D. popularis* has enlarged third and fought tarsi segments and fifth segment is silvery-white (fig. 1).

*D. pennatus* and *D. subpennatus* is almost indistinguishable in most morphological characters using in the keys of *Dolichopus* species. *D. subpennatus* was assigned as a separate species from a series of *D. pennatus* specimens according to the structure of the male cilioratum (tibial organ) [D’assis-Fonseca, 1976].

Ornamented segments of the middle tarsi are presented in four out of five species, while in *D. lineatocornis* males the tarsi are simple. Females of these species are similar morphologically (cryptic), their distinctive features are of a comparative nature. Difficulties chiefly arise due to broad intraspecific morphological variation between females of the *D. pennatus* group, which can seriously confound their identification. Females of *D. popularis* are distinguished in the group by two apical bristles on hind femora and mainly yellow antennae, while the color of the antennae in other species ranges a lightly according to geographical distribution of population: postpedicel is always completely black, pedicel varies from dark with a narrow yellow base to dark on top and yellow on the inner and ventral sides. *D. lineatocornis* females differ from *D. pennatus* females by a slightly more yellow base of the first segment of the middle tarsi. *D. argyrotarsis* females are absent from the existing identification tables.

Differences between females of *D. pennatus* and *D. subpennatus* were presented in the species description and keys [D’assis-Fonseca, 1976, 1978]. *D. pennatus* share the following characters: metanotun share small spines, scutellum covered with more or less dense hairs. Hairs located only at the lower edge of the scutellum and bare metanotum are characterized for both sexes of *D. subpennatus*. However, the further morphological analysis of specimens from Urals and Western Siberia showed that the characters is variable and cannot be used in the species diagnosis of females [Selivanova et al., 2019]. In addition, the species can be distinguished from each other by the shape of the apicoventral epandrial lobe [Selivanova et al., 2019]. The use of the listed characters requires experienced entomologists for correct identification or availability of collection material for comparison, therefore, the diagnosis of these species is most often carried out according to the modified middle legs of males.

## Materials and Methods

Landmark-based geometric morphometric analysis and the methods of traditional morphometry were used to evaluate of the interspecific differences. Wing shape and legs morphometric characters variation was observed from 236 specimens of the *D. pennatus* group from Voronezh State University (Russia) insects collection: 22 females and 56 males of *D. argyrotarsis*, 12 females and 24 males of *D. lineaticornis*, 40 females and 42 males of *D. pennatus*, 18 females and 8 males of *D. popularis*, 8 males and 6 females of *D. subpennatus*. The wings and legs of each specimen was removed, transfer to a microscope slide and covered with the cover slip. Each wing was photographed at 20 magnification using a digital camera Levenhuk C NG attached to a stereomicroscope.

### Geometric morphometric analysis of wing shape

To describe wing shape nine homologous Type 1 landmarks, placed in the vein junctions and vein terminations, were used (fig. 2). Each landmark have been digitized using TpsDig-2.32 software [Rohlf, 2008].

**Fig. 2.**
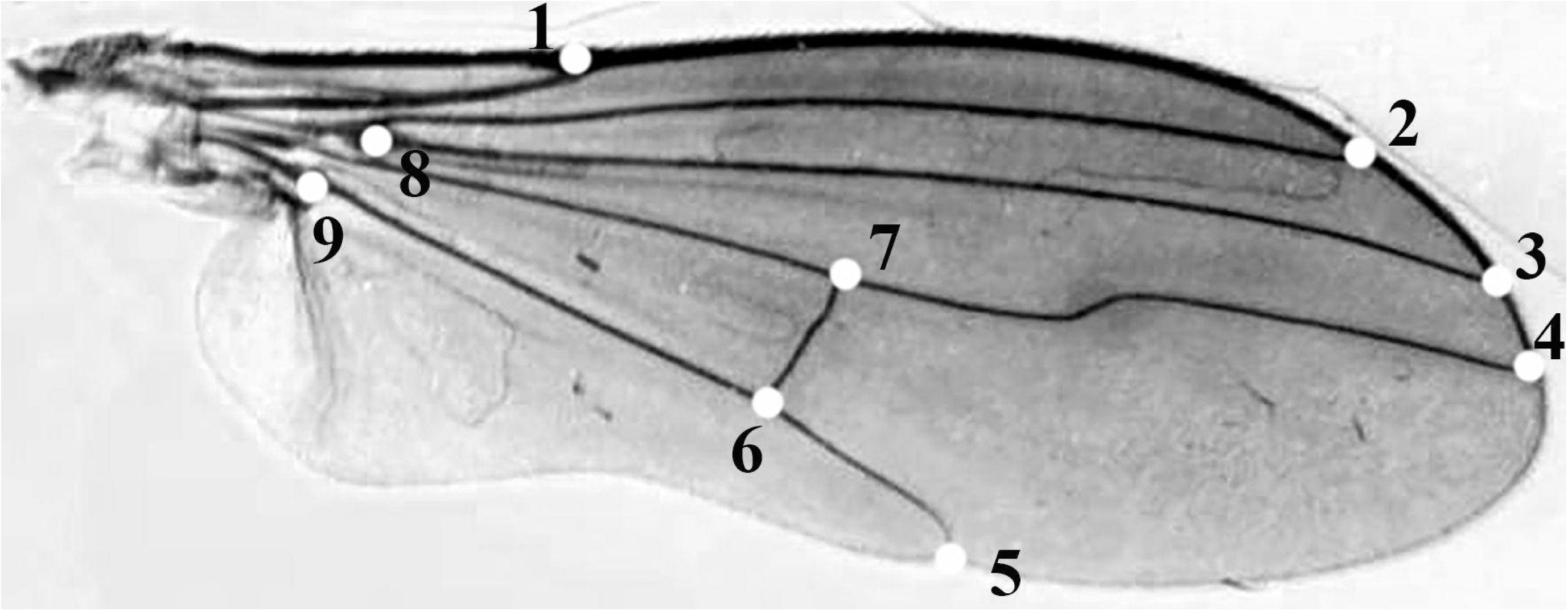
Positions of the landmarks used for studying wing shape. (*D. argirotarsis*, male).

Centroid size (square root of the sum of squared distance between each landmark and the wing centroid) was used as the measure of size for each wing. Procrustes coordinates obtained from landmark data were used for further statistical analyses of wing shape. For this purpose, all wings were superimposed and differences in the position of the landmark points were then examined. The superimposition was done with the Procrustes technique (wings are scaled to a unit centroid size, superimposed on an origin so that their mean x and y coordinates becomes (0, 0), the landmarks are finally rotated so that the distance between all the landmarks of all specimens and a consensus configuration becomes minimal) [Rohlf, 1999]. Then, analysis was carried out using the methods of multivariate statistical analysis in software MorpholJ [Klingenberg, 2011] и Statistica for Windows (version 10).

Tests for significant differences among species were undertaken using a one-way ANOVA. Canonical variate analysis (CVA) was used to determine the most important characters as a possible discriminator between species and evaluate species relative distribution in shape space. To quantify the differences between the wing shapes of the species, the Procrustean distance was calculated [Zelditch, Swiderski, 2004]. The significance of the differences was tested using a permutation test with

### Morphometric analysis of legs

Measurements were made by the photos of the specimens using the ImageJ software (1.53b) [Schindelin et al., 2015]. Nine morphometric characters of the legs were measured: the lengths of the fore-, mid-, and hind femora (F1, F2, and F3), the fore-, mid-, and hind tibia (T1, T2, and T3), and the first segment of the fore-, mid-, and hind-tarsi (tar1, tar2, and tar3). Then the following twelve relative signs were calculated: the ratio of the lengths F1 to T1, F1 to tar1, F1 to F2, F1 to F3, T1 to tar1, T1 to T2, T1 to T3, F2 to T2, F2 to tar2, T2 to tar2, F3 to T3, and F3 to tar3.

Test for significant differences among species were undertaken using an analysis of variance (ANOVA). To examine relationship among species based on legs morphometric characters, unweighted pair group method with arithmetic average (UPGMA) was performed using PAST 3.09 software [Hammer, 2001].

### Molecular data analysis

The analyzed molecular matrix included molecular sequences of the mitochondrial gene encoding cytochrome c oxidase (COI) (810 characters). The study included both sequences previously deposited in GenBank [GenBank, 2021] and sequences carried out especially for this study by the Sintol Enterprise (Russia). In total molecular sequences of 25 species was studies. Amplification and sequencing were performed using the methods and primers described in previous studies [Bernasconi et al., 2007a, b].

Phylogenetic tree was constructed using the maximum likelihood method in MEGA software (Kumar et al., 2018). The significance of the inner branching pattern was estimated by a bootstrap analysis with 1000 pseudo-replicates. As a measure of phylogenetic signal of legs morphometric characters, we used Pagel’s lambda (λ) [Pagel, 1999] and Blomberg K-statistic [Freckleton et al., 2002]. A Pagel’s lambda is an indicator that can take a value from zero to one, and a value close to one indicates a more significant phylogenetic signal in character. To calculate Pagel’s lambda, the *phylosyg* function *phytools* package [Revell, 2012] was used in R environment [R Development Core Team, 2014]. Blomberg K-statistic also takes values from zero to one, but if the phylogenetic signal is very high, then K-statistic can rise over one. To calculate Blomberg K-statistic, the *Kkalk* function *picante* package was used in R environment [Kembel et al., 2010]. For testing purpose the indications of differences of the metric from 0, a p-value was obtained by randomizing the trait data 999 times.

## Results

Since the sexual dimorphism was shown to be valuable in the morphometric characters of the wings legs [Chursina, Negrobov, 2018a, b], males and females were analyzed separately. Results of ANOVA exhibit highly significant differences among males of the species both in wing centroid size: F = 5.86, P = 0.0001, and in wing shape: Wilks’ lambda = 0.027; F = 27.0; df = 56, 974.62; P < 0.0001. Procrustes distances between average wing shapes for males also were significant (P < 0.0001). The greatest distance – 0.0687 и 0.0620 were found, respectively, between *D. argyrotarsis* – *D. popularis* and *D. lineaticornis* – *D. popularis*, the smallest distance – 0.0138 – between *D. argyrotarsis* and *D. subpennatus*.

The first canonical variate (CV1) explain 64.10% of the overall wing shape variation. The main differences in wing shape described by CV1 occur in the position of the posterior transverse vein (*dm-m*) and the apical segment of *M*_*4*_ and the shape of the wing apex (fig. 3). The first axis described a separation *D. popularis* from *D. pennatus* and the group of species including *D. argyrotarsis, D. lineaticornis, D. subpennatus*. The second canonical variate (CV2) included 21.78% of the total shape dispersion. CV2 clearly separate specimens of *D. popularis*. It should be noted that males of *D. argyrotarsis, D. lineaticornis* and *D. subpennatus* are weakly differentiated by wing shape.

**Fig. 3.**
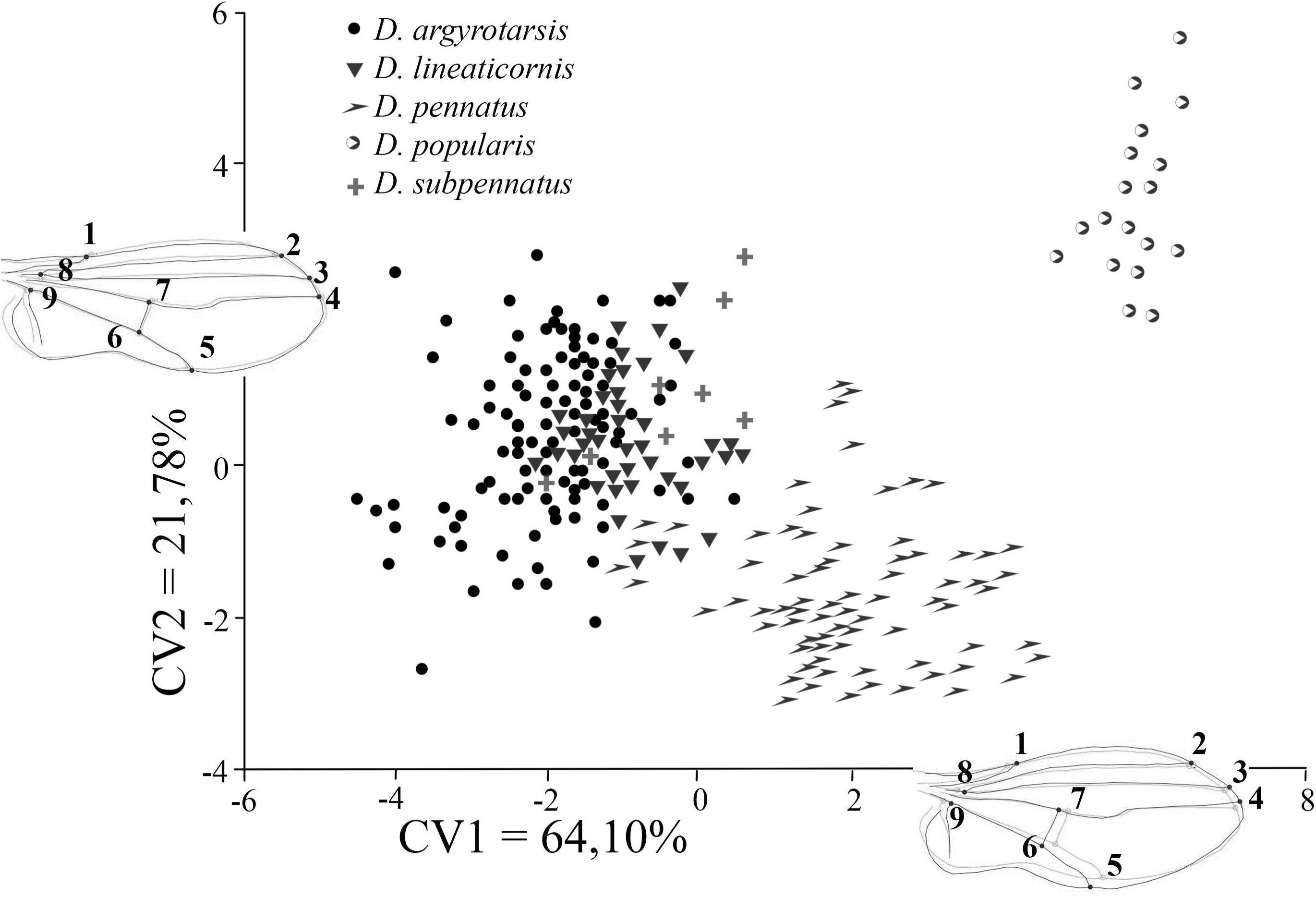
Scatter plot from CVA showing scores of the first two CVs for male of the five species of *Dolichopus* with shape changing schemes.

We observed a high significant differences between females in both centroid wing size: F = 37.8, P < 0.0001, and wing shape: Wilks’ lambda = 0.0506; F = 13.8; df = 56, 671; P < 0.0001. The first canonical variate (CV1) included 48.62% of the total wing shape variation. The main differences in the female wing shape described by CV1, as in males, concerned changes in the area of Landmarks 5 and 7 (the locations of *dm-m* and the apical part of *M*_*4*_) (fig. 3). Females of *D. popularis* are reliably distinguished along the CV1 axis. An important role in the separation of species was also played by the second canonical variable (CV2 = 40.51%), which described the proximal displacement of the *dm-m* and reliably separated *D. argyrotarsis* females from females of four other species. Females of *D. lineaticornis, D. pennatus* and *D. subpennatus* in wing shape cannot be differed.

The main differences in the shape of the wing between species in females concerned the same landmarks as in males, which is confirmed by the low value of the angle between the first (36.0°, P = 0.0003) and the second (45.1°, P = 0.003) main principle components.

The ANOVA of morphometric leg characters also showed significant differences between males: Wilks’ lambda = 0.168; F = 5.0; df = 48, 433; P < 0.0001, and females: Wilks’ lambda = 0.192; F = 3.0; df = 48, 287; P < 0.0001.

The UPGMA dendrogram analysis (fig. 5) revealed that males of the studied species are more similar to each other than to females of their species. The leg morphometric characters of *D. argyrotarsis* male with modified middle tarsi were more similar to those of *D. lineaticornis* males, rather than species that also have legs ornaments. According to the leg morphometric characteristics, the males of *D. popularis, D. pennatus*, and *D. subpennatus* with high bootstrap support were combined into one group.

**Fig. 4.**
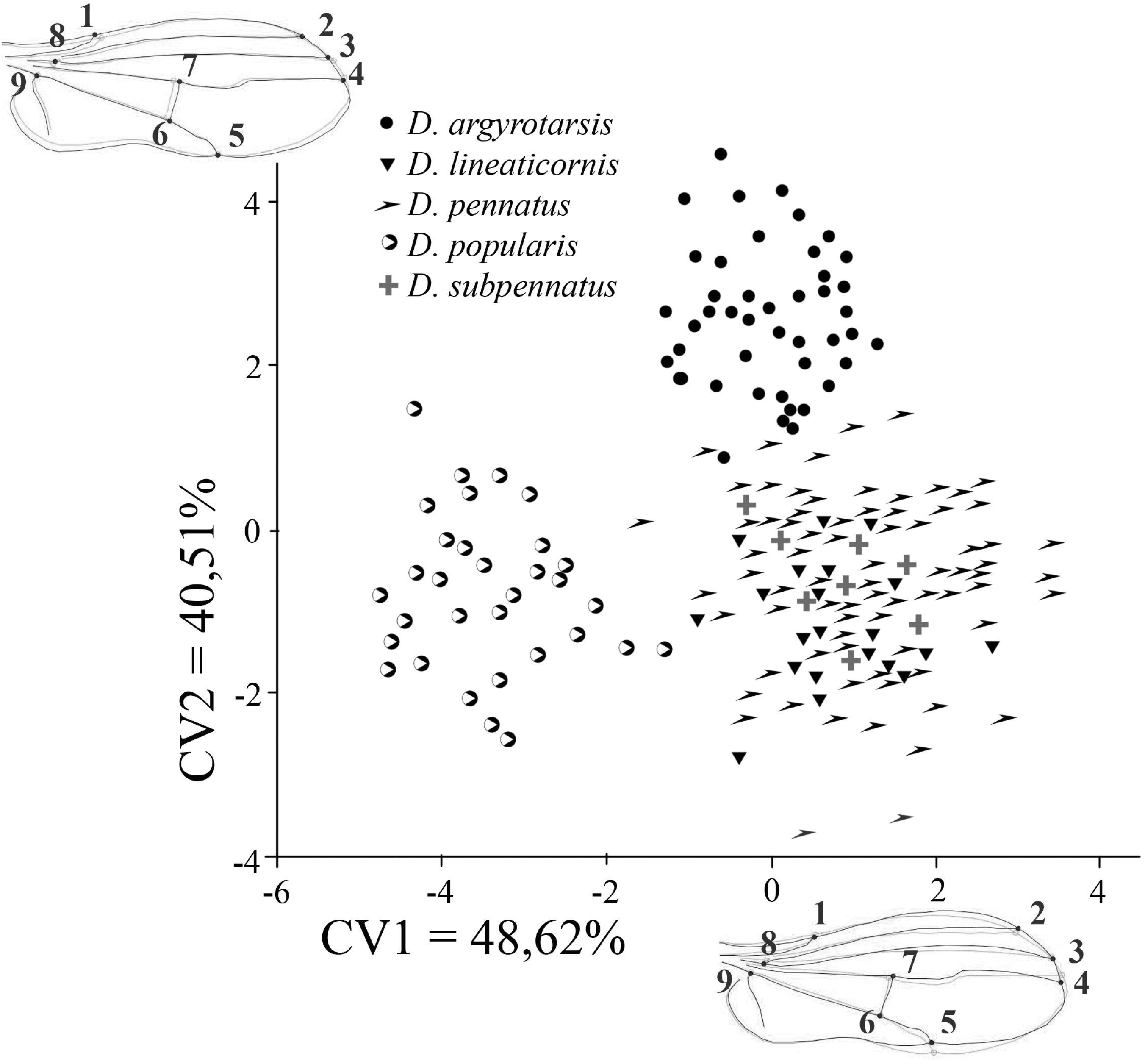
Scatter plot from CVA showing scores of the first two CVs for female of the five species of *Dolichopus* with shape changing schemes.

**Fig. 5.**
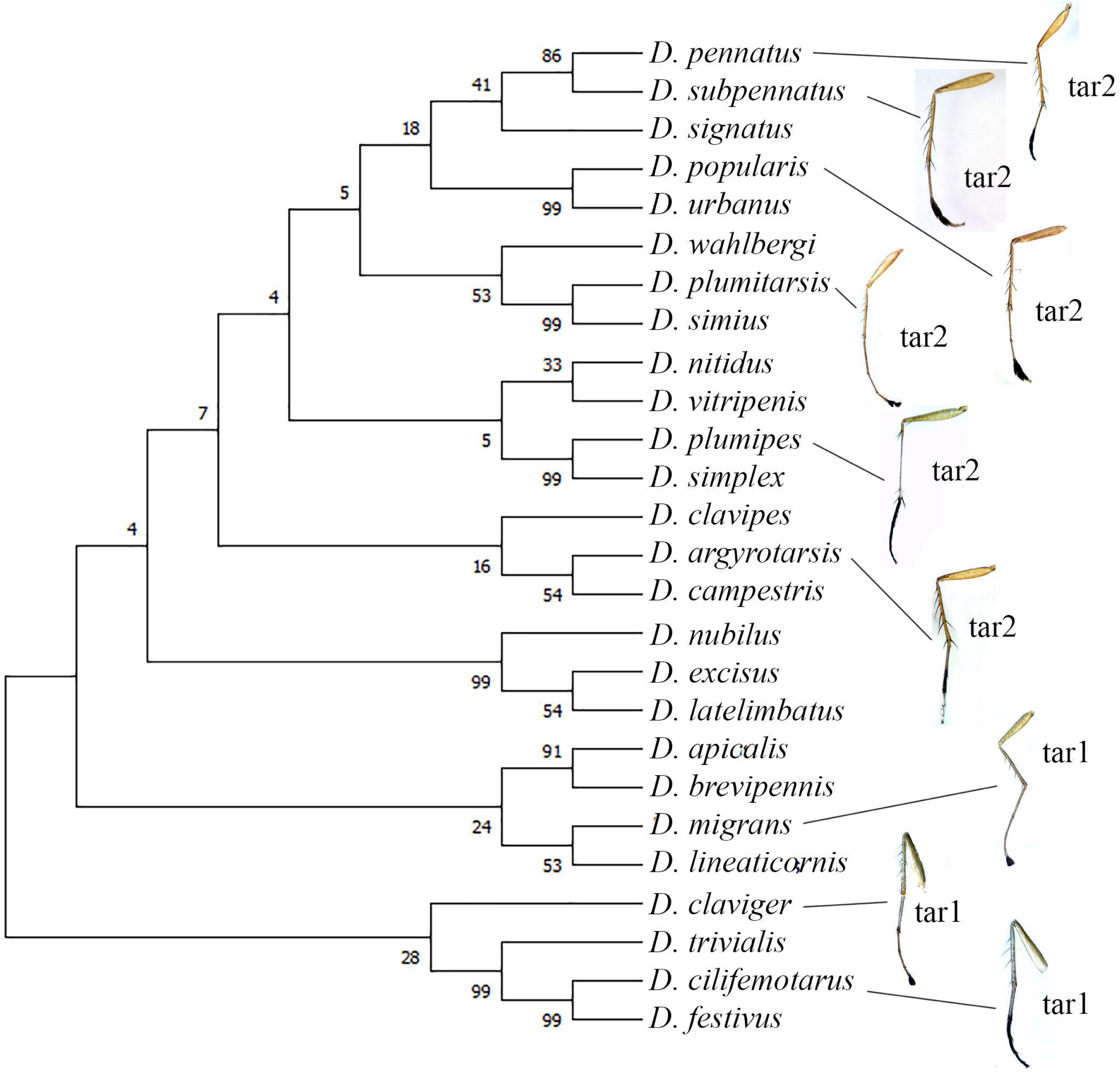
Results of UPGMA cluster analysis of legs morphometric characters of the five *Dolichopus* species.

The main differences in the legs morphometric characters between males lie in the relative length of the first segment of middle legs (tar2). Males of *D. pennatus, D. subpennatus* and *D. popularis* have significantly longer tar2 than males of *D. argyrotarsis* and *D. lineaticornis*, which is most evident in the F2/tar2 ratio (P < 0.0001): in *D. popularis* males this ratio is 1.54, *D. subpennatus* – 1.55, *D. pennatus* – 1.56, *D. argyrotarsis* and *D. lineaticornis* – 1.71. When comparing the morphometric features of the legs of *D. subpennatus* and *D. pennatus*, no significant differences were found.

Differences in legs morphometry in females were less statistically significant. *D. argyrotarsis* females differed from females of the other four species in the shortened first segment of the hind legs, which is reflected in the ratio F3/tar3 (P < 0.01): in *D. popularis* females, this ratio is 2.09, in *D. pennatus, D. subpennatus* and *D. lineaticornis* females – 2.11, in *D. argyrotarsis* females – 2.27.

Significant differences between *D. subpennatus* and *D. pennatus* females were showef by the F2/tar2 ratio (P = 0.002), which was 1.81 and 2.07, respectively, that is, *D. subpennatus* females had a relatively shorter first segment of the middle legs. Other discovered characters are presented in table 2.

**Table 1.** Procrustes distances between average wing shape for males (over the main diagonal) and females (under the main diagonal) of five *Dolichopus* species.

|  | *D. argyrotarsis* | *D. lineaticornis* | *D. pennatus* | *D. popularis* | *D. subpennatus* |
| --- | --- | --- | --- | --- | --- |
| *D. argyrotarsis* |  | 0,0209<br>P < 0,0001 | 0,0129<br>P < 0,0001 | 0,0200<br>P = 0,005 | 0,0163<br>P = 0,002 |
| *D. lineaticornis* | 0,0172,<br>P < 0,0001 |  | 0,0141<br>P = 0,0001 | 0,0212<br>P = 0,036 | 0,0116<br>P = 0,22 |
| *D. pennatus* | 0,0348<br>P < 0,0001 | 0,0316<br>P < 0,0001 |  | 0,0195<br>P = 0,0006 | 0,0099<br>P = 0,41 |
| *D. popularis* | 0,0687<br>P < 0,0001 | 0,0620<br>P < 0,0001 | 0,0387<br>P = 0,0005 |  | 0,0189<br>P = 0,37 |
| *D. subpennatus* | 0,0138<br>P = 0,013 | 0,0232<br>P < 0,0001 | 0,0288<br>P < 0,0001 | 0,0612<br>P < 0,0001 |  |

**Table 2.**
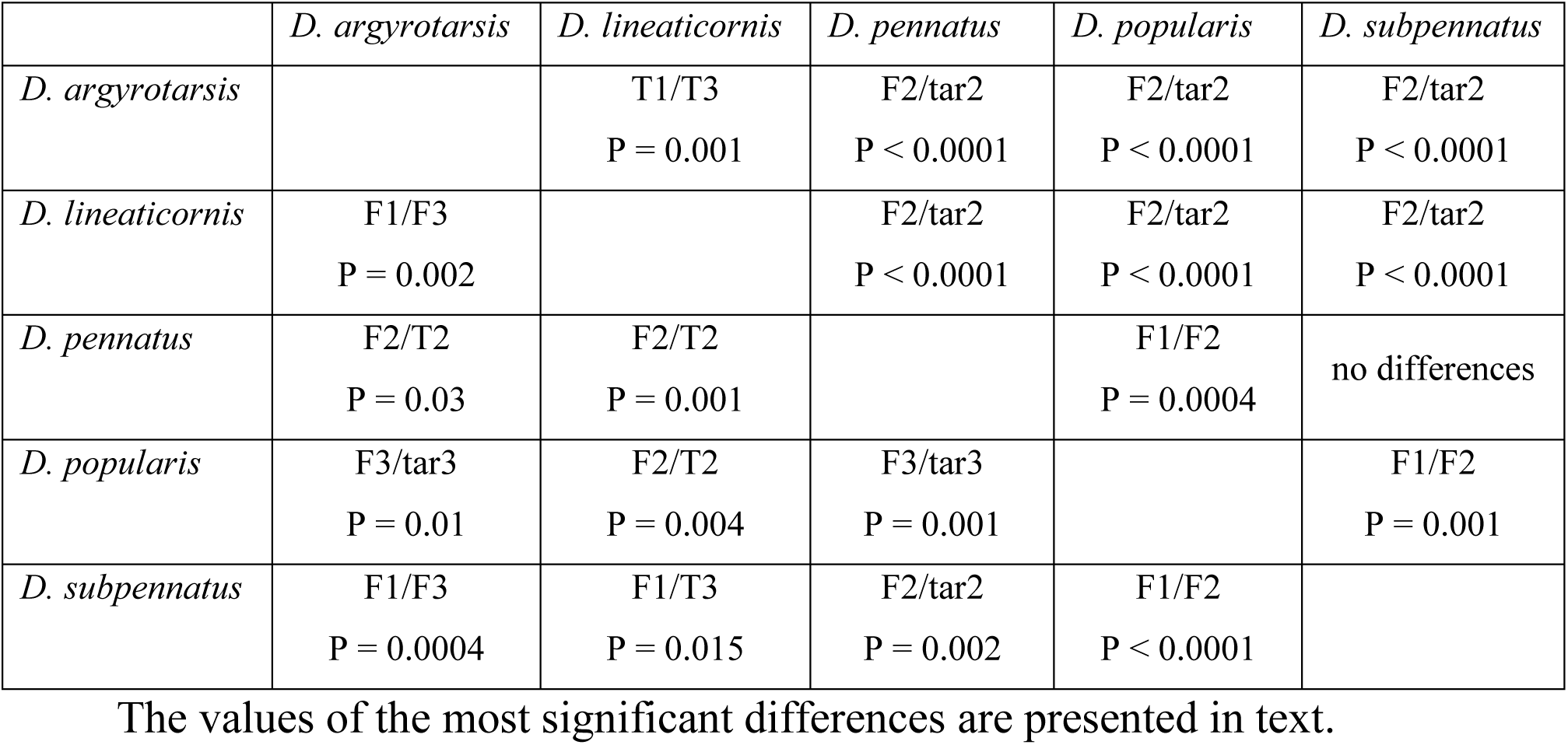
Discovered characters of leg morphometry for male (over the main diagonal) and females (under the main diagonal) of five *Dolichopus* species.

Analysis of molecular data showed that, despite morphological similarity, not all of these species are forming one phylogenetic clade (fig. 6). For instance, both species *D. pennatus* and *D. subpennatus* (bootstrap support value was 86) have modified middle legs; both species *D. popularis* and D. urbanus (99) have modified middle legs. Whereas in the pair of such species as *D. argyrotarsis* and *D. campestris* (54) – male *D. campestris* are characterized by simple legs, but male *D. argyrotarsis* is characterized by ornamented middle tarsi. In the pair of such species as *D. lineaticornis* and *D. migrans* (53) – the middle tarsi are ornamented in the first species, and the fore tarsi are ornamented in the second species. According to the calculated parameters, ornamented of fore legs have significant phylogenetic signal. Pagel’s lambda was 0.99, P < 0.00001 and Blomberg’s K was 1.098. This suggests that the enlargement of the forelegs (*D. simius, D. plumitarsis, D. claviger*) or the presence of erect hairs on fore tarsi (*D. cilifemoratus, D. festivus, D. trivialis*) occurred in phylogenetically close species of *Dolichopus*.

**Fig. 6.** Maximum likelihood tree, obtained from COI sequence of 26 *Dolichopus* species with illustrations of ornaments of the fore (tar1) and middle (tar2) legs of males.

At the same time, in groups with high bootstrap support, a species with ornamented legs and a species with simple legs were combined together: for example, *D. plumipes* and *D. simplex* (bootstrap support value was 99), *D. apicalis* and *D. brevipennis* (99). For such character as the ornamented middle legs, Pagel’s Lambda was 0.48, P = 0.27, Blomberg’s K – 0.202, that is, the phylogenetic signal of the character is absent in the analyzed species group. Thus, parallel evolution in ornamented middle legs is quite likely.

## Discussion

All five studied species together with the very rare *D. signatus* Meigen, 1824, which was not considered in this study, according to identification key of A.A. Stackelberg [1933] were included in the *Eudolichopus* group, distinguished as a subgenus; that the subgenus status was canceled [Steyskal, 1973]. Further, for the convenience of using the identification key, only conditional group III was distinguished, characterized by yellow legs and yellow postocular cilia in the lower part of the head. This group includes two-thirds of the Dolichopus species with varying degrees of similarity in a number of other characters.

The “*pennatus*” species group has a number of common morphological characters. The following characters are common to the five species: the presence of only one anterodorsal seta on the middle tibia, the absence of erect hairs, one preapical seta on the middle and hind femora, the absence of a dorsal seta on the first segment of the middle legs, 2–3 dorsal setae on the first segment of the hind tarsi. The following color features also are characteristic of these species: yellow scape and black pedicel and postpedicel, calypter yellowish with black squamal cilia; thorax and abdomen are metallic green and yellow fore coxae and dark middle and hind coxae. The characteristic characters for this species group is a smoothly curved medial vein and a pronounced anal wing lobe.

Such feature of sexual dimorphism as ornamented legs is often seen in the Dolichopodidae family. According to published data, males use ornamented legs in courtship rituals [Sivinski, 1997]. Similar patterns of sexual behavior could arise independently in different taxonomic groups and lead to the formation of a similar combination of morphological characters [Bonduriansky, 2006; Chursina, 2019]. It is critical to examine such characters is important for construction of robust phylogenetic hypothesis, since in the case of convergent evolution, such structures do not reflect the phylogenetic closeness of species.

According to the data obtained as a result of comparative analysis of morphological and molecular characters, a sufficiently high phylogenetic signal in *Dolichopus* species is present in the ornaments of fore leg, while the phylogenetic signal of the middle leg ornaments is low. In other words, modifications of the middle legs occur in not phylogenetic closely species and in most cases may indicate homoplasy.

Analysis of the wings and legs morphometric characters demonstrated that in the studied species, similar leg ornaments were associated with the formation of a certain habitus: *D. pennatus, D. popularis* and *D. subpennatus* males in addition to the extended last segments of the middle tarsus, were also characterized by an elongated first segment of the same tarsi, a shortened wing with a proximally displaced *dm-m*.

This study is the first to quantify subtle wing and leg morphometric traits among species of the “*pennatus*” species group. Although the morphology of male legs has been widely used in identification of the species, diagnostics of females was difficult. However, based on wing shape and legs morphometric characters presented here we found significant differences between some of the studied species. *D. pennatus* and *D. subpennatus* females can be discriminating with female of *D. popularis* and *D. argyrotarsis* using wing shape analysis, although no significant differences were found with *D. lineaticornis*. It is proposed to distinguish *D. argyrotarsis* females from *D. pennatus* and *D. subpennatus* by the shorter first segment of the middle tarsi.

## Acknowledgements

The work was funded by RFBR and NSFC according to the research project No 20-54-53005.

